# Genomic sequencing of Lowe syndrome trios reveal a mechanism for the heterogeneity of neurodevelopmental phenotypes

**DOI:** 10.1101/2021.06.22.449382

**Authors:** Husayn Ahmed Pallikonda, Pramod Singh, Rajan Thakur, Aastha Kumari, Harini Krishnan, Ron George Philip, Anil Vasudevan, Padinjat Raghu

**Author notes:** Present Address: Cancer Dynamics Laboratory, The Francis Crick Institute, London, UK. Equal contribution (Husayn Ahmed Pallikonda, Pramod Singh).

## Abstract

Lowe syndrome is an X-linked recessive monogenic disorder resulting from mutations in the *OCRL* gene that encodes a phosphatidylinositol 4,5 bisphosphate 5-phosphatase. The disease affects three organs-the kidney, brain and eye and clinically manifests as proximal renal tubule dysfunction, neurodevelopmental delay and congenital cataract. Although Lowe syndrome is a monogenic disorder, there is considerable heterogeneity in clinical presentation; some individuals show primarily renal symptoms with minimal neurodevelopmental impact whereas others show neurodevelopmental defect with minimal renal symptoms. However, the molecular and cellular mechanisms underlying this clinical heterogeneity remain unknown. Here we analyze a Lowe syndrome family in whom affected members show clinical heterogeneity with respect to the neurodevelopmental phenotype despite carrying an identical mutation in the *OCRL* gene. Genome sequencing and variant analysis in this family identified a large number of damaging variants in each patient. Using novel analytical pipelines and segregation analysis we prioritize variants uniquely present in the patient with the severe neurodevelopmental phenotype compared to those with milder clinical features. The identity of genes carrying such variants underscore the role of additional gene products enriched in the brain or highly expressed during brain development that may be determinants of the neurodevelopmental phenotype in Lowe syndrome. We also identify a heterozygous variant in *CEP290*, previously implicated in ciliopathies that underscores the potential role of *OCRL* in regulating ciliary function that may impact brain development. More generally, our findings demonstrate analytic approaches to identify high-confidence genetic variants that could underpin the phenotypic heterogeneity observed in monogenic disorders.

## Introduction

The oculo-cerebro renal syndrome of Lowe, commonly referred to as Lowe syndrome (LS) is an X-linked recessive disorder characterized by a triad including a developmental cataract, intellectual disability and altered renal function. LS is a rare disease with a reported prevalence of 1:100000 and about 400 cases are reported in the literature [reviewed in (Staiano et al., 2015)]. Affected children are born with a range of clinical features including congenital cataracts, infantile glaucoma, neonatal or infantile hypotonia, intellectual impairment and renal tubular dysfunction. LS is believed to be a monogenic disorders; the defective gene has been identified, denoted *OCRL* and encodes an inositol polyphosphate 5‐phosphatase (Attree et al., 1992) *OCRL* encodes a multidomain protein; in addition to the core 5-phosphatase domain, the protein also contains multiple other domains including an N-terminal PH domain, an ASH domain and a Rho-GAP domain that are proposed to control its localization and protein-protein interactions. *OCRL* is part of a large family of ten inositol polyphosphate 5-phosphatases; based on the domain structure present along with the 5-phosphatase domain, 5 sub-families are defined; *OCRL* and the related gene *INPP5B* form one of these subfamilies (Ooms et al., 2009). The *OCRL* transcript is widely expressed across human tissue and cell types and it is thus intriguing that mutations in this gene only impact the eye, brain and kidney as assessed by clinical findings. At the level of individual cells, the OCRL protein is reported to localize to multiple organelle membranes including the plasma membrane, Golgi apparatus and the endo-lysosomal compartment [reviewed in (Mehta et al., 2014)]. As a result, OCRL is proposed to regulate a large number of membrane transport activities as well as the cellular cytoskeleton.

Despite being a monogenic disorder arising from mutations in the OCRL gene, the range of clinical features reported in LS patients is variable. Although the ocular, cerebral and renal findings are involved in most cases, the extent to which each of these three organs is affected can be quite variable. For example, some patients with LS show very mild intellectual disability whereas in other individuals, the mental retardation is substantially more severe. Conversely in some individuals only the renal phenotypes are reported; this condition, also known as Type II Dent’s disease is also a consequence of mutations in the OCRL gene. Despite the analysis of more than 400 individual patients, a mechanism to explain this clinical variability in the setting of this monogenic disorder has not been put forward. In this study we present the genetic analysis of a family affected by LS. Using the interesting genetic architecture of this family coupled with whole genome sequencing we analyzed the coding exome sequence of each of the affected patients in this family. Our findings indicate that additional variants in genes affecting neurodevelopment contribute to the variable neurodevelopmental phenotype seen in the individual members of this family. Our findings also suggest a genetic framework within which one may predict disease severity and progression and hence planning of clinical management in patients with LS.

## Results

### Clinical characterization

We analyzed a family identified through a male proband born to non-consanguineous parents, who presented to the clinic at 7 months of age with a history of delayed developmental milestones. Clinical analysis revealed a history of bilateral congenital cataract (corrected surgically), global developmental delay and hypotonia. Parents noticed polyuria and polydipsia few months prior to presentation. Based on the clinical triad of neurodevelopmental delay, altered renal function and congenital cataract in a young male child, a tentative diagnosis of LS was made. Detailed laboratory investigations excluded other potential etiologies such as endocrine disorders, inherited muscle disorders and a TORCH panel were negative ruling out an infectious etiology. Physical examination revealed features of rickets with laboratory investigations revealing normal anion gap hyperchloremic metabolic acidosis, normal renal functions and laboratory features of vitamin D deficiency. Urine routine microscopy revealed nephrotic range proteinuria without glycosuria or aminoaciduria along with phosphaturia and hypercalciuria. Ultrasonography revealed normal kidney structure. A diagnosis of LS was made based on clinical features and laboratory findings which was confirmed by genetic testing. The child was initiated on oral bicarbonate and potassium supplements along with enalapril along with early intervention program for his developmental delay. A brain MRI performed at 2 years of age showed bilateral symmetrical hyperintense lesions regions in T2 weighted images in frontal and parietal regions suggestive of periventricular leucomalacia. There was no diffusion restriction in FLAIR images or ventriculomegaly or atrophy. At the last follow up at 9 years of age, the proband demonstrated improvement in most domains of development except in speech. He could walk without support, showed improvement in hyperactivity, social skills and in school performance as well with mental age of 4 years, but his speech was limited to monosyllables only even at 9 years of age. Brain auditory evoked potentials were normal. Renal function was normal (eGFR 96 ml/min/1.73m^2^), although proteinuria and hypercalciuria persisted.

We documented a full family history of the proband (Fig 1 A); this revealed two maternal cousins, identical twin boys with a similar triad of congenital cataract, renal dysfunction and neurodevelopmental delay, although the extent of the renal and neurodevelopmental delay were variable. In the case of the twin cousins, the neurodevelopmental delay seemed less severe than in their single cousin with speech delayed developing at 2 and 4 years. A formal developmental assessment at age 7 revealed stereotypic behavior although both children were able to attend regular school with a satisfactory academic performance. Renal investigations revealed nephrotic range proteinuria, normal anion gap metabolic acidosis, hypercalciuria without glycosuria or aminoaciduria, hypophosphatemia and high serum alkaline phosphatase. Renal function was normal. Treatment was initiated on oral bicarbonate and potassium supplements along with ACE inhibitor enalapril. All three children showed growth retardation with height z-scores in the range of −2 to 4. All three also developed bilateral cataracts which was corrected by a lensectomy at an early age.

**Figure 1:**
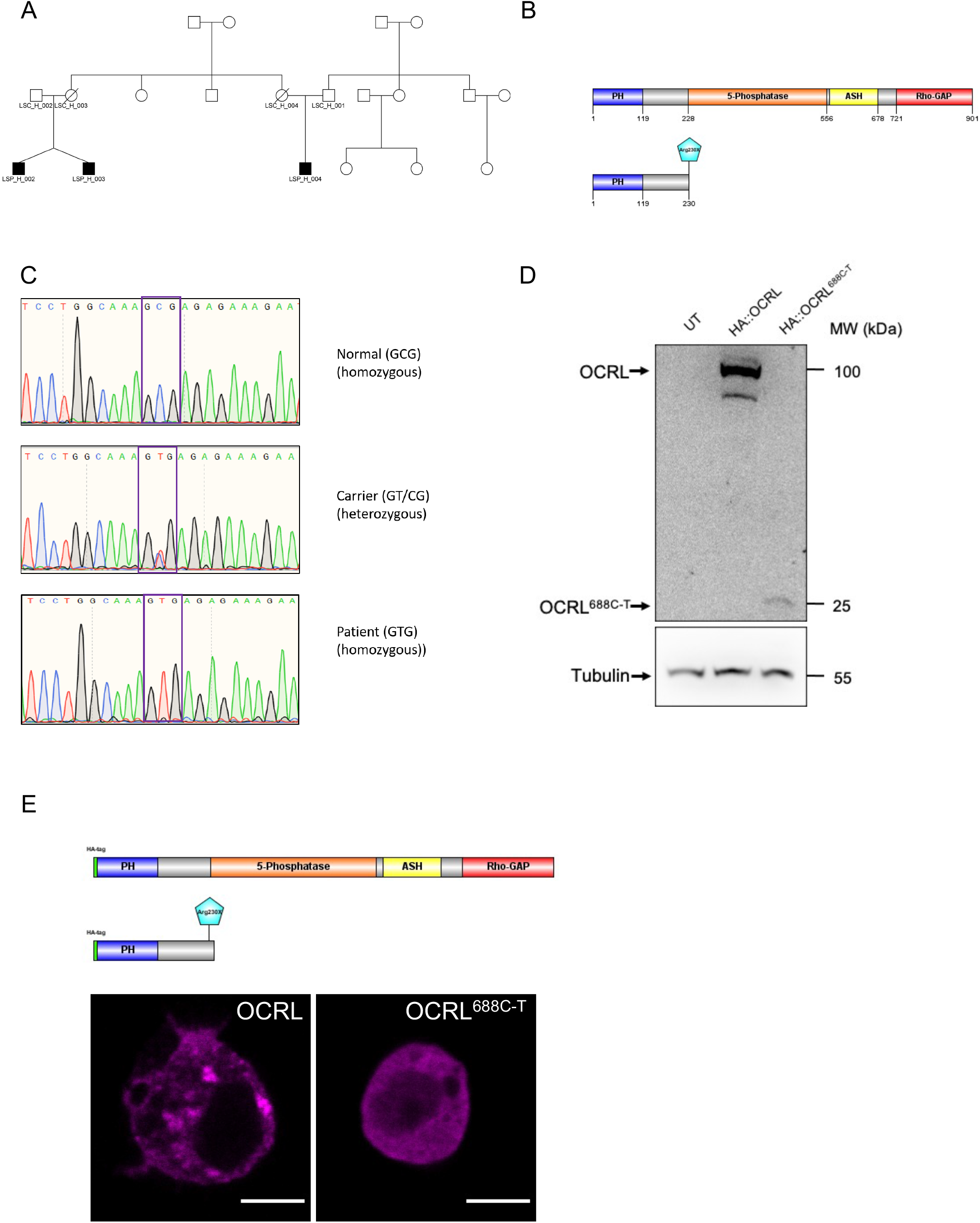
(A) Pedigree of the family analyzed in this study. Affected individuals are showed as shaded symbols. (B) Domain structure of the wild type OCRL gene is shown. PH, 5-phosphatase, ASH and Rho-GAP domain are all shown. The predicted protein produced by the wild type and patient cDNA is depicted. (C) Sanger sequencing chromatogram showing the GCG to GTG nucleotide change within exon 8 of the OCRL gene. (D) Western blot analysis of wild type and patient cDNA for OCRL transfected into *Drosophila* S2R+ cells expressing Actin-GAL4. UTC-untransfected control, pUAS-HA::hOCRL (wild type) and pUAS-HA::hOCRL^688C-T^ (patient). The truncated band produced by the patient cDNA is shown. Tubulin is used as a loading control. (E) protein localization of HA::hOCRL and HA::hOCRL^688C-T^ transfected into *Drosophila* S2R+ cells and visualized using immunolabelling. Schematic of the HA tagged proteins from wild type and patient cDNA is shown. Magenta fluorescence represents the localization of the protein. Write scale bar is 5μm

### Genetic analysis

Given the clinical triad of developmental delay, altered renal function and cataract along with a potential pattern of X-linked inheritance (mothers unaffected), strongly suggests an X-linked recessive disorder consistent with a diagnosis of LS. In order to confirm the diagnosis of LS, we performed Sanger sequencing of DNA extracted from peripheral blood. We sequenced all 23 exons of the OCRL gene form all three patients and their parents. In each of the three patients we identified the same single base pair change- c.688C>T; this change was also seen but in only one allele in each of the maternal genomes (Fig 1C). The c.688C>T mutation described introduces a p.Arg230X in the 8^th^ exon and is predicted to truncate the protein just prior to the lipid phosphatase domain of OCRL (Fig 1B). The variant c.688 C>T in OCRL meets the criteria for the following - PVS1, PM2 and PP3 and is thus categorized as Pathogenic according to the American College of Medical Genetics and Genomics (ACMG) classification (Richards et al., 2015). These findings provide genetic evidence that along with clinical findings confirm the diagnosis of LS in all three patients; the proband and his two cousins.

To confirm the impact of the described mutation, we expressed either wild type OCRL and the c.688C>T variant in *Drosophila* S2R+ cells. In the case of the wild type cDNA, this resulted in the expression of a protein with the expected Mr of 110 kD; in the case of the c.688C>T variant, a truncated protein of ca. Mr 25 kD was produced (Fig 1D). We also looked at the localization of the proteins; the wild type protein showed a widespread punctate distribution throughout the cell whereas the c.688C>T variant showed a diffuse distribution throughout the cytosol (Fig 1E).

### Whole genome and exome sequencing

In order to understand the genetic landscape within which mutations in OCRL result in clinical phenotypes, we performed next generation sequencing on blood derived DNA of all three affected children and their parents; both whole exome sequencing (WES) and whole genome sequencing (WGS) was performed. Overall, we obtained a uniform depth of coverage in all samples sequenced in both WES and WGS (Fig 2A, B) and the vast majority of sequences could be aligned with the human reference genome sequence (Fig 2C,D). For each sample, post alignment processing was performed (Fig 2G) and variant calling was done using the GATK Haplotype caller (McKenna et al., 2010). For the purpose of Trio analysis joint genotyping was done using a consolidated VCF file and the data were processed using the pipeline depicted in Fig 2G. Using this approach, a list of variants was identified for each sample in both WES and WGS experiments. This resulted in an equivalent load of variants in each sample; as expected, the vast majority of these were single nucleotide polymorphisms (SNPs) with a smaller number of deletions and insertions in each sample (Fig 2 E,F).

**Figure 2:**
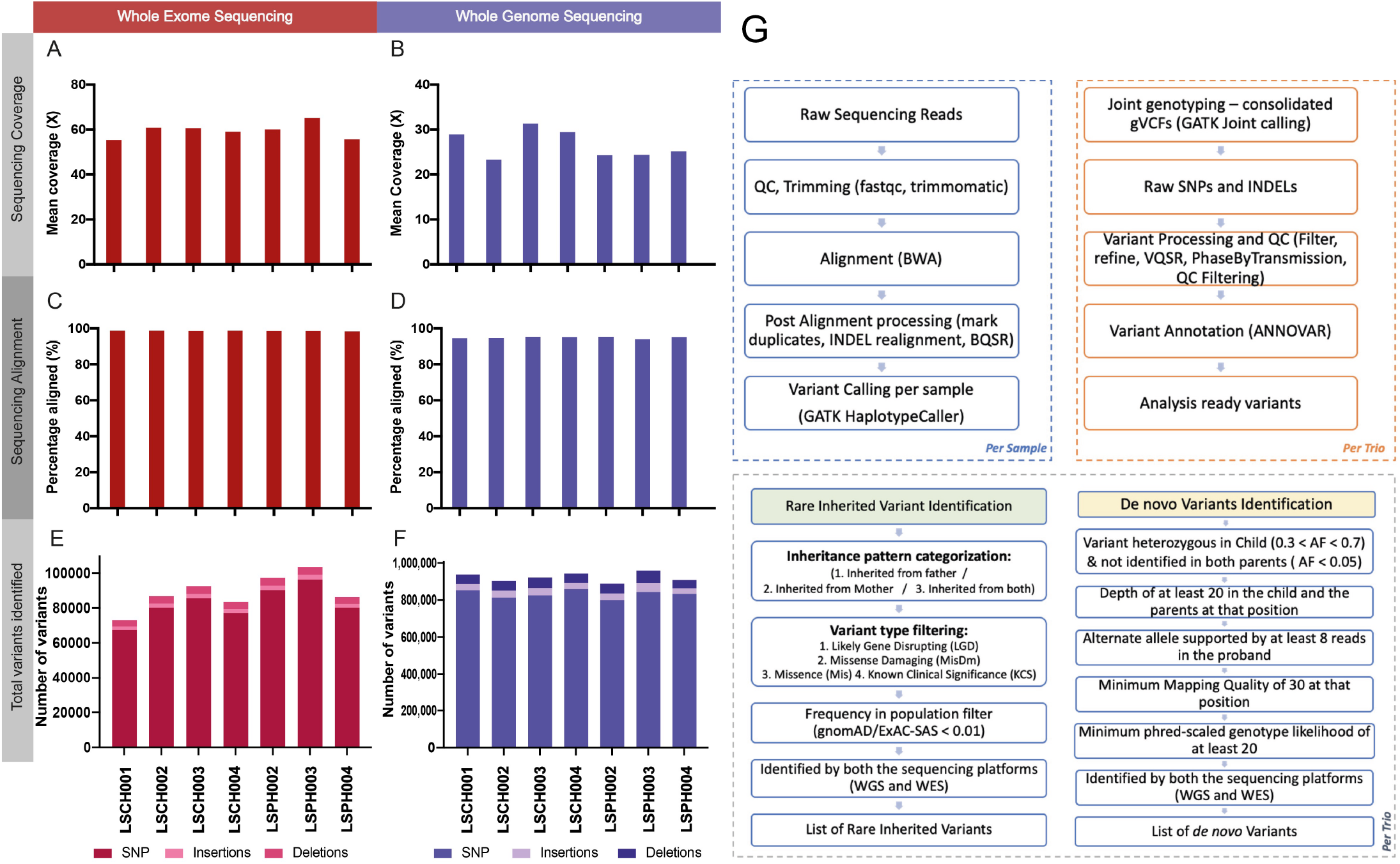
Summary of whole genome and exome sequencing and variant discovery. (A) Mean sequencing coverage in whole exome sequencing (WES) and in (B) whole genome sequencing (WGS) for all samples is shown. (C) The percentage of reads that aligned to the reference genome hg19 in WES and (D) in WGS. (E) A stacked bar plot showing the number of SNPs, insertions and deletions identified from WES and (F) WGS (G). Overview of the WGS/WES analysis and variant prioritization pipeline. The data processing workflow that was utilised to identify rare inherited and de novo variants is shown here. See methods for more details on this workflow. BQSR: Base Quality Score Recalibration, gVCF: Genomic Variant Call File, VQSR: Variant Quality Score Recalibration, ExAC-SAS: Exome Aggregation Consortium South Asian Cohort, AF: Alternate Allele Frequency,

### A strategy to evaluate variants that could contribute to LS phenotypes

A current challenge in genome sequence analysis in the context of human disease is to estimate the functional importance of the large number of variants that are described when comparing any genome with a reference genome sequence. To prioritize variants of likely functional importance in the context of LS, we defined a three-tiered system (3TS) for categorization of candidate genes likely to be important in contributing to the LS phenotype and attempted to shortlist variants in this defined set of genes. Earlier sequencing studies on genetic disorders have adopted a similar tiered approach to assess and identify the most likely candidate genes and have been successful in doing so (Taylor et al., 2015). We defined Tier 1 genes as those which are known to cause disorders, abnormalities or phenotypes that are observed in or very similar to that of the LS and those genes which are known to physically interact with the OCRL protein. The Tier 2 genes were defined as those that are involved in the biological processes in which the OCRL protein is proposed to play an important role. Tier 3 genes were defined as those which are the direct interactors of Tier 1 genes (See methods).

The Tier 1 genes were further categorized into four - Ocular (genes causal of ocular phenotypes), Cerebral (genes causal of cerebral phenotypes), Renal (genes causal of renal phenotypes) and OCRL Interactome (genes interacting directly with OCRL) categories (Fig 3B). This tiered system of categorizing enabled a systematic approach to understand the mutational burden in any individual with different levels of relevance and significance. For example, a variant in any candidate gene within these tiers is likely to be more relevant to the phenotype compared to variants in other genes. Further, a variant in a Tier 1 gene is likely to have higher relevance or significance compared to that of Tier 2 and tier 3. Of the 22,287 protein coding genes in the human genome (Salzberg, 2018), using this approach, we restricted our analysis to variants in 15077 genes within tiers 1, 2 and 3. Of these 2106 genes were in Tier 1 with the largest proportion being in Tier 3 (Fig 3A). The prioritized rare inherited and de novo variants were annotated with this 3-tiered system of candidate gene categorization in order to evaluate the functional importance of the newly identified variants. A full list of genes in each of the tiers in provided in Supplementary table 3.

**Figure 3:**
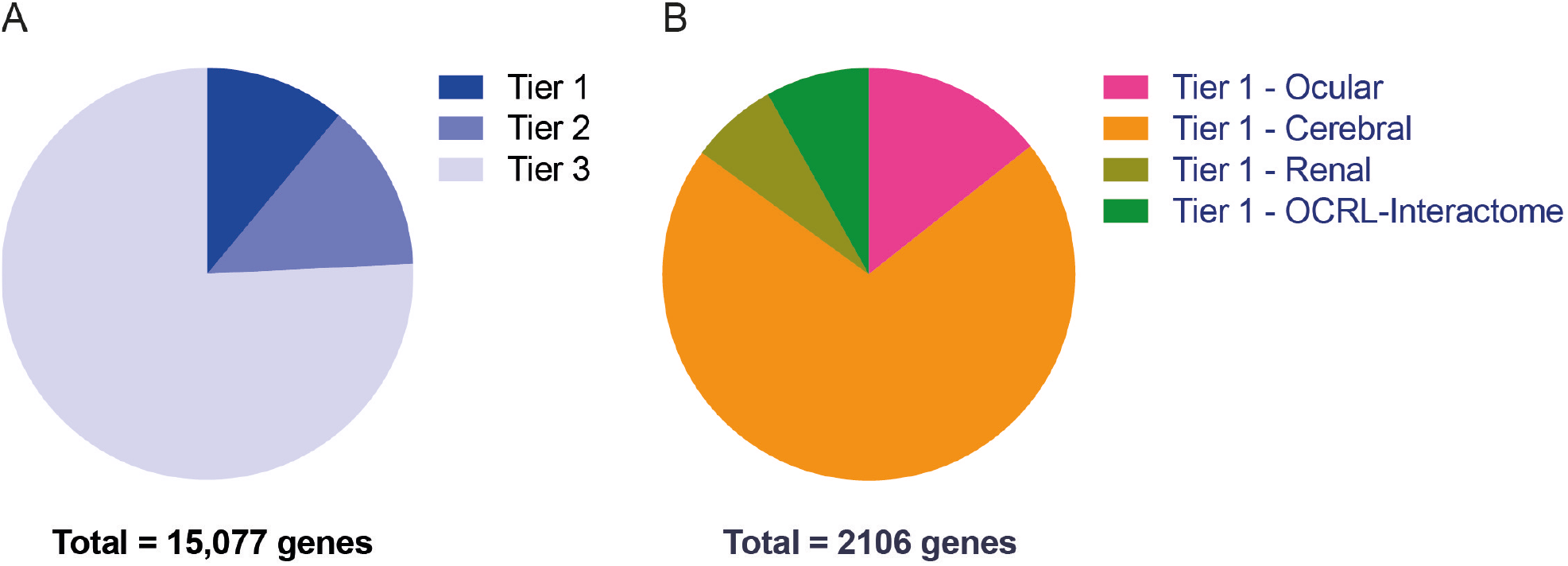
Summary of 3-Tiered System of categorization of candidate genes. (A) A pie chart showing the distribution of the number of genes in each tiers of the 3-Tiered system of categorization of candidate genes. Please see the methods for the details on curation of these tiered gene lists. (B) The distribution of four sub-categories of genes in tier 1 is shown as a pie chart.

### Rare inherited variants in LS

In order to discover rare inherited variants, we devised a pipeline that filtered the total variant set in each individual to establish the inheritance of each variant from one or both parents (Supplementary Figure 1). All such inherited variants were then assessed to determine that they belonged to one of the following categories (i) Likely gene disrupting (LGD) i.e introducing a premature termination codon or altering the reading frame of the protein (ii) Missense (Mis)-SNPs that change the amino acid residue in a protein (Fig 4A) (iii) Missense Damaging (MisDm) Those Mis variants assessed by *in silico* variant effect predicting algorithms as likely to alter the structure and therefore function of the protein (Fig 4B) (iv) variants previously determined to be of known clinical significance (KCS). Variants of all these categories were classified as rare if their frequency in the population of sequenced genomes (gnomAD/ExAC-SAS) was <0.01. Lastly, we used only those variants that were identified in both the WES and WGS experiments for any given individual. Using these criteria, we shortlisted 19 LGD variants in the affected children (Fig 4C). Each of the children carried a different number of LGD variants; LSPH002 (12), LSPH003 (9) and LSPH004 (8) with the twins P2 and P3 carrying 8 identical variants. Of this set of 19 LGD variants, 15 (78.9%) were in genes represented in any of the 3 tiers of our defined 3TS geneset (Figure 3A). One variant, in the gene CEP290 belonged to tier 1 (cerebral and renal sub-categories). There were 3 (15.8%) LGD variants identified in tier 2 genes and 14 (73.7%) LGD variants identified in tier 3 genes. Of these LGD variants 2 were found in all three boys, 6 were found in the identical twins LSPH002 and LSPH003 but not in LSPH004 (Fig 4D). Conversely, there were 6 variants that were found only in LSPH004 but not in the identical twins LSPH002 and LSPH003.

**Figure 4:**
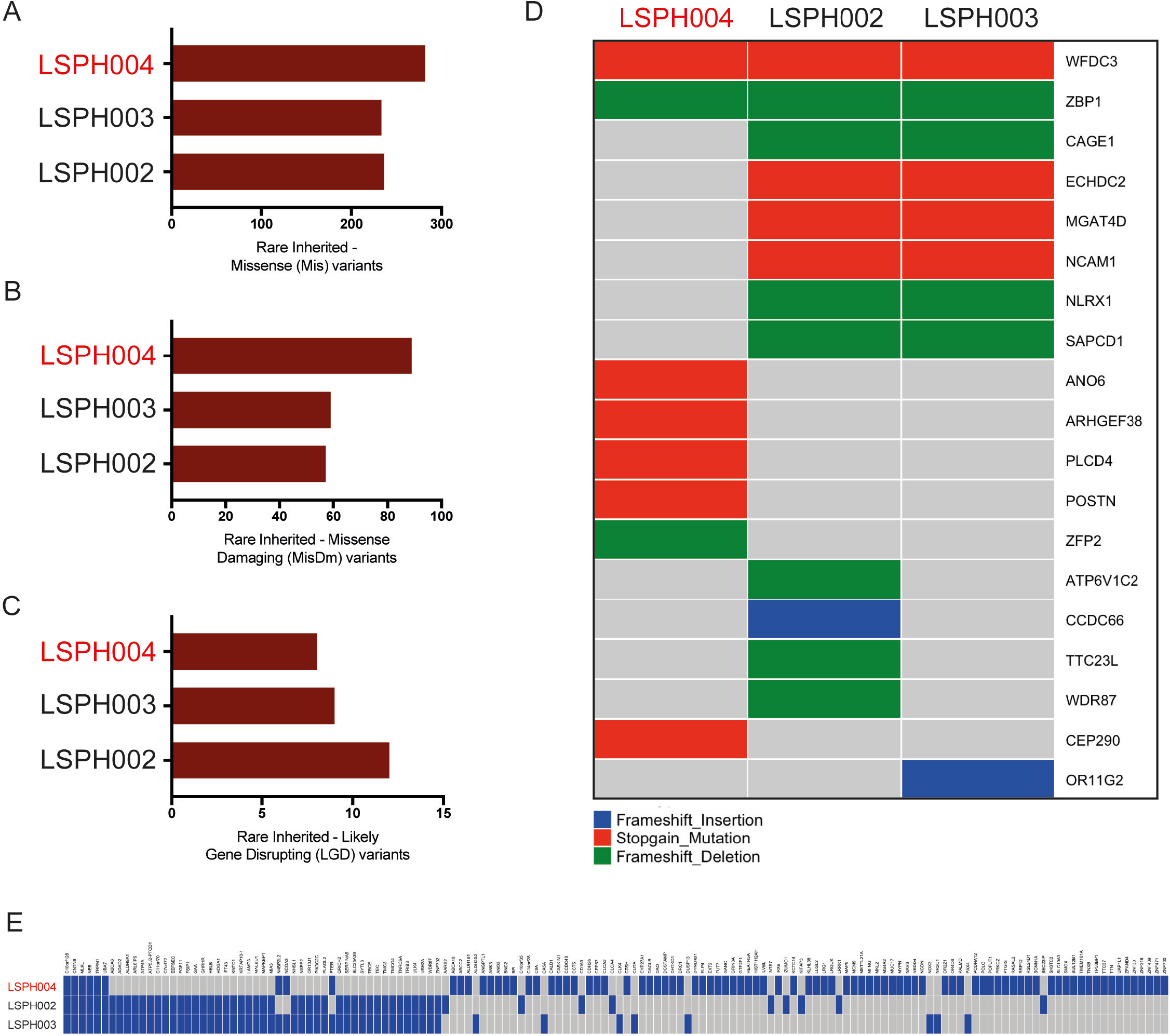
Prioritized rare inherited variants. The variants identified in the OCRL patients were subjected to prioritization for identifying the most functionally relevant variants. The prioritization approach identified rare inherited variants from these patients that are functionally relevant to LS. (A) Total number of rare inherited missense (B) Inherited missense damaging and (C) Inherited likely gene disrupting variants in each patient is shown. (D) A summary of prioritized rare inherited variants that are categorized as Likely Gene Disrupting (LGD) and the genes that carry these variants are summarized here. The presence of stopgain variants (coloured red), frameshift insertions (blue) and deletions (green) across the 3 LS patients are shown. (E) Similarly. the summary of rare inherited variants categorized as Missense Damaging (MisDm) is shown. A complete list of prioritized variants with their gene annotation is provided in the Supplementary Table 1.

Apart from the LGD variants, we identified 146 missense damaging (MisDm) variants. While 9 (6.2 %) of these variants were identified in the tier 1 genes, 13 (8.9 %) and 86 (58.9 %) were identified in tier 2 and tier 3 genes respectively (Fig 4E). Of these only 6 were found in all three children; 82 were found only in LSPH004 and 41 were found in both LSPH002 and LSPH003 but not in LSPH004. A complete list of prioritized variants with their gene annotation is provided in the Supplementary Table 1.

### *De novo* variants in OCRL trios

Earlier sequencing based studies of trios have shown the contribution of de novo variants and in many cases their higher impact than inherited variants in neurological disorders (Liu et al., 2018), neurodevelopmental disorders (McRae et al., 2017; Wright et al., 2015), schizophrenia (Fromer et al., 2014) and autism spectrum disorders (Iossifov et al., 2014). In families with developmentally normal parents, whole exome sequencing of the child and both parents resulted in a 10-fold reduction in the number of potential causal variants that needed clinical evaluation compared to sequencing only the child. In this study, we tried to identify and assess the contribution of *de novo* variants in LS.

In that context, we employed the sequence data for identification of DNMs. An unfiltered set of putative DNMs is highly enriched for errors (Malhotra et al., 2011; Sebat et al., 2007). Hence, we employed methodologies and recommendations from earlier studies which reported identification and validation of DNMs in large-scale trio datasets (Fromer et al., 2014; Jiang et al., 2013; Sanders et al., 2012)(McRae et al., 2017; Willsey et al., 2017) to reduce errors. *De novo* variants were identified as heterozygous variants in a child that were not present in either of the parents. In addition, we employed a number of quality control criteria [Fig 2G and materials & methods) to reduce the false positive rate of *de novo* variant reporting. This resulted in the discovery of a limited number of *de novo* variants in all three children (Fig 5A). As expected, for *de novo* variants, we found minimal overlap between the variants identified in the identical twins LSPH002 and LSPH003 and the ones found in LSP004 were also different (Fig 5B). Of these only one, in C5 was present in LSPH002 and LSPH003 but not in LSPH004 whereas the converse was true of one other variant in ARSD.

**Figure 5:**
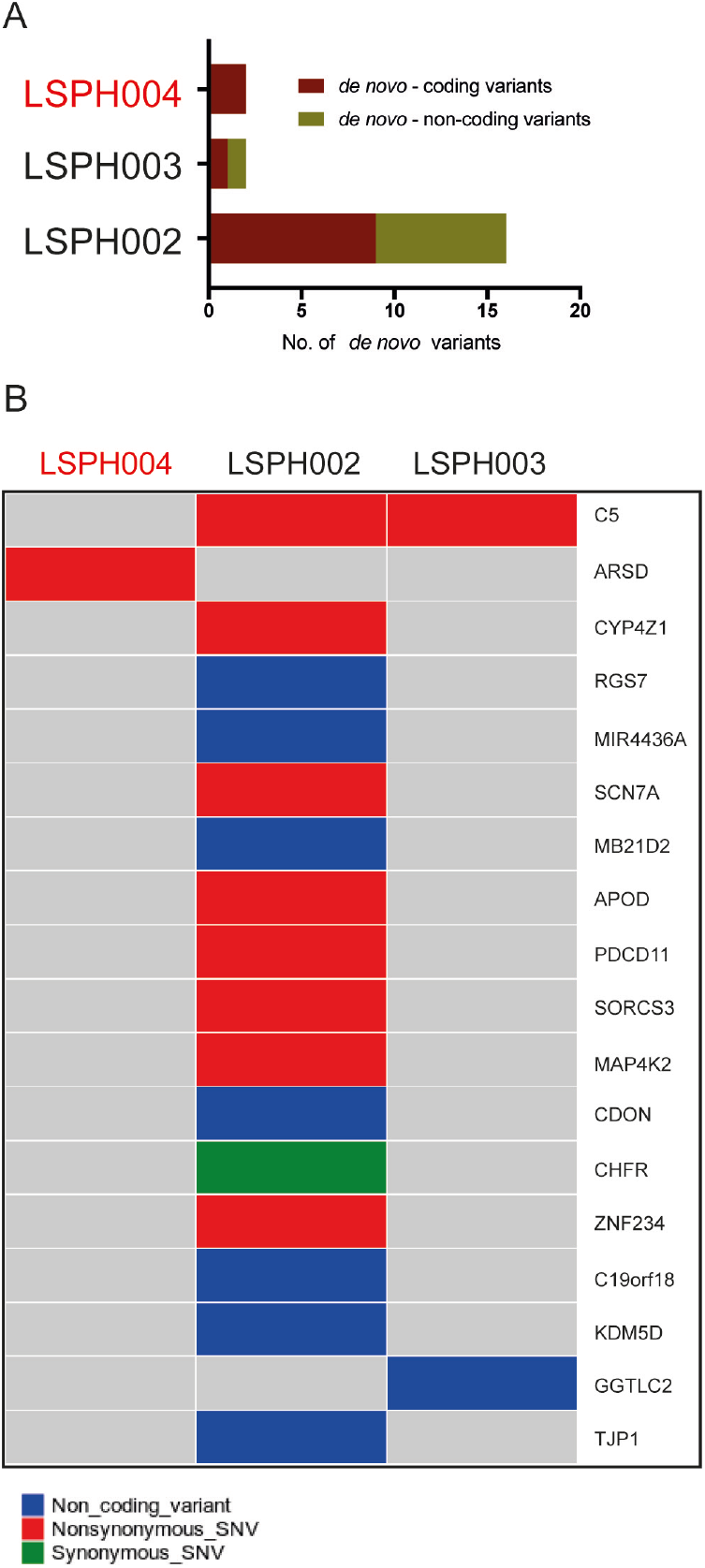
Prioritized *de novo* and rare inherited variants. (A) Total number of prioritized *de novo* variants identified in each probands is shown. Coding and non-coding *de novo* variants are coloured differently. (B) Prioritized *de novo* variants and the genes that carry these variants are summarized here. They are coloured by the type of the variant. Non-synonymous (red), synonymous (green) and non-coding (blue) *de novo* variants across the 3 LS patients are shown.

## Discussion

Although monogenic disorders are defined on the basis of a single gene whose altered function results in a clinical phenotype, it is recognized that there can be a considerable degree of variability in the spectrum and severity of clinical features between individual patients with monogenic disease such as thalassemia (Weatherall, 2001) and cystic fibrosis (Shanthikumar et al., 2019). Such phenotypic variability has also been noted in the case of neurodevelopmental disorders such as Rett syndrome (Neul et al., 2010; Neul et al., 2019). A number of factors can contribute to such phenotypic variability including the nature of the mutant allele in a specific patient, environmental factors, variable allelic expression (e.g X-inactivation in the case of X-linked genes) and also variants in other genes that alter their function and hence impact phenotype. Given that LS is an X-linked recessive disorder, variable X-linked inactivation is unlikely to contribute to disease variability in affected male patients. In this study, we analyzed three patients, from a single extended family, all of whom have the identical mutation in OCRL that produces a truncated protein without the phosphatase domain. Despite this, phenotypic variability was seen between these three patients, with a clear dichotomy in the neurodevelopmental phenotype between the identical twins and their single cousin. Thus, the nature of the mutant allele and it’s biochemical impact cannot explain the phenotypic variability observed in this set of LS patients. A large number of mutations distributed across the length of the OCRL protein have been described in LS patients [summarized in (Staiano et al., 2015)]. Amongst these are a number of alleles in whom the position of the stop codon truncates the protein prior to the core phosphatase domain, effectively generating functionally null alleles; analysis of clinical phenotypes in patients carrying such mutant alleles reveals wide variations between these patients (Supplementary Table 3). These observations are consistent with our finding that the nature of the mutant allele in OCRL1 is not sufficient to account for phenotypic variability between patients.

A likely reason for the variable phenotypes in individual LS patients is the impact of modifier variants in the genome, where genes containing such variants are likely to impact the function of cells lacking OCRL. Genetic modifiers play an important role in human development influencing the relationship of phenotype and genotype (Slavotinek and Biesecker, 2003); however the role of such background genetic variation in the clinical heterogeneity of LS has not been examined. In this study we carried out a comprehensive analysis for such variants in three children from a single family all of whom carry the same truncating mutation in the OCRL gene. Using a combination of WES and WGS, we identified a set of rare inherited and *de novo* protein coding variants present in each child, that could contribute to the variability of the LS neurodevelopmental phenotype. This analysis revealed a set of 88 such variants present only in LSPH004 but not in LSPH002 and LSPH003. Such variants could in principle act as enhancers of the effect of OCRL depletion in the brain and therefore contribute to the enhanced neurodevelopmental phenotype of this patient. We analyzed the expression of genes carrying these variants in the human brain using the GTEx database (Aguet et al., 2017) and found that only a subset of these genes (Fig 6 A,B) showed significant expression in the brain; In many cases (Fig 6A) high levels of expression in the brain were seen during pre-natal development with a dynamic pattern of gene expression noted during the various stages of prenatal development. Given that the neurodevelopmental phenotypes of LS are evident soon after birth, the activity of genes that which show such patterns of elevated and dynamic gene expression during the pre-natal period are likely to be part of the molecular and cellular mechanisms processes leading to normal brain development. The altered activity of these gene products may therefore be important in terms of the cognitive and neurodevelopmental phenotypes.

**Figure 6:**
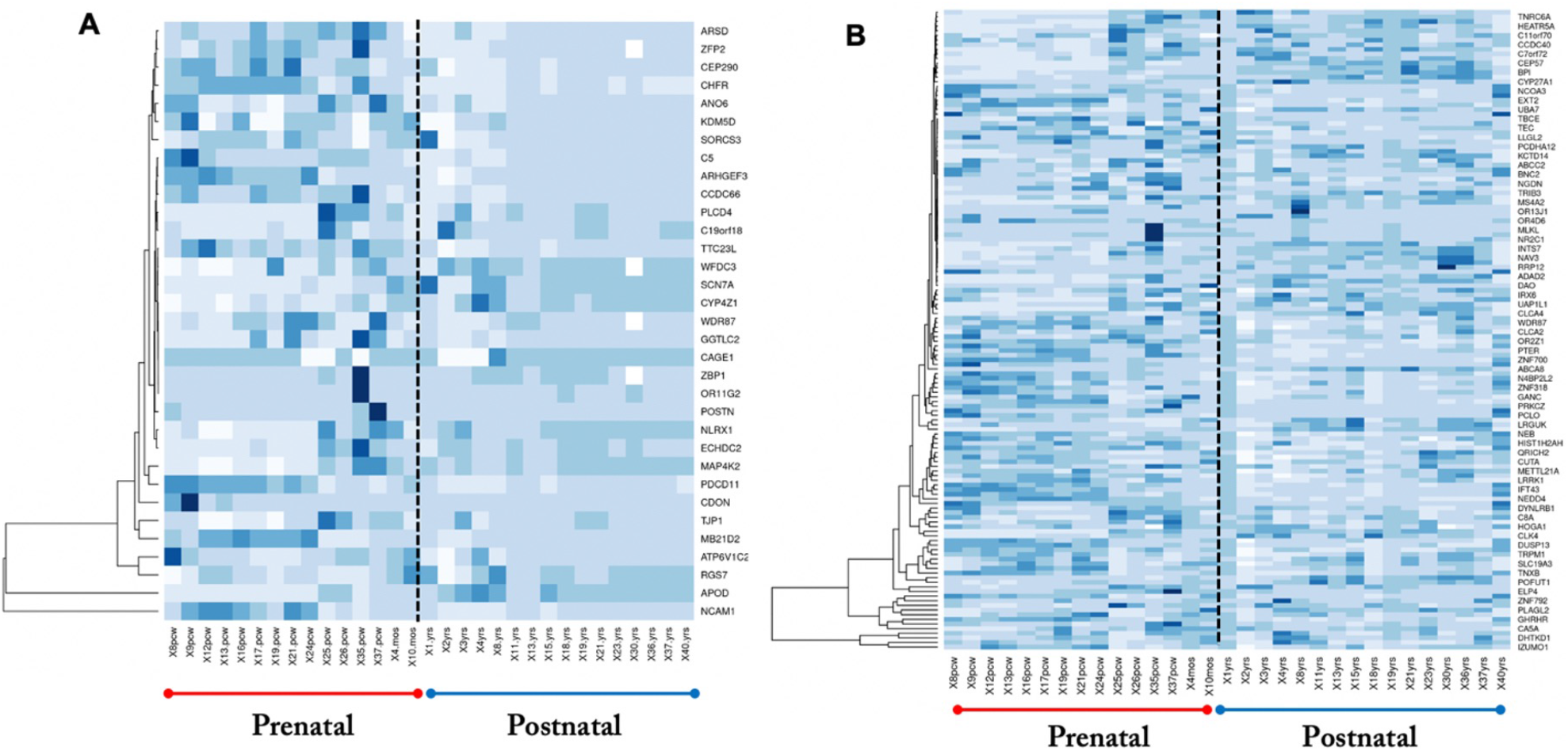
Brain expression pattern of genes with (A) LGD and (B) MisDm variants identified in the three patients with LS in this study. X-axis shows the age in weeks post-conception (pcw) and years post birth (yrs). Individual genes are listed along the Y-axis. Level of expression is color coded as: Dark blue-high expression, White-low expression.

Among the variants that we found was a heterozygous stop gain variant identified in the tier 1 gene CEP290 (p.G1890X). While this variant is carried by both the mothers, it was found to be maternally inherited only by the single child LSPH004. The mutational landscape of CEP290 has been studied in the context of ciliopathies (Coppieters et al., 2010). Homozygous and compound heterozygous variants in CEP290 have earlier been reported as pathogenic in various ciliopathies, including Nephronophthisis, Joubert syndrome, Meckel syndrome, Senior-Loken syndrome, Leber congenital amaurosis and Bardet-Biedl syndrome. The role of CEP290 in ciliogenesis has been widely studied and is also known to be involved in the ciliary transport processes, regulation of the ciliary membrane composition and ATF4-mediated transcription (Shimada et al., 2017; Wu et al., 2020). Although CEP290 has never been reported in the context of LS, it is intriguing to note that mutations in INPP5E a member of the same 5’phosphatase superfamily as OCRL have been associated with Joubert syndrome itself a ciliopathy (Bielas et al., 2009; Jacoby et al., 2009; Travaglini et al., 2013) and recent studies indicated an important role for phosphoinositides in ciliary biology (Conduit and Vanhaesebroeck, 2020). Given the altered neural development noted in many ciliopathies (Suciu and Caspary, 2021), it is likely that the variant in CEP290 that we have identified in LSPH004 may contribute to the more severe neural phenotype of this individual. CEP290 is expressed at high levels during in utero development in the fetal brain and also shows a dynamic expression pattern (Fig 6A), supporting a likely role for this variant in the disease phenotype of LSPH004.

Another stopgain heterozygous variant was identified in the gene PLCD4 (tier 2 and tier 3 gene, p.Q632X) in the single child LSPH004 as maternally inherited. PLCD4 belongs to the family of phosphoinositide specific phospholipases and hydrolyses phosphatidylinositol 4,5-bisphosphate (PIP2) to mediate intracellular calcium signalling. A recent study has proposed an unexpected partnership of a *Drosophila* phospholipase C (dPLCXD) and PTEN that impacts the accumulation of PI(4,5)P2 on endosomes, and compensates for the loss of the 5-phosphatase OCRL in *Drosophila* cells (Mondin et al., 2019). Mondin et al., also showed that treatment with a PLC activator m-3M3FBS reduces the consequences of the loss of OCRL, also in the absence of PTEN, thereby suggesting a therapeutic strategy for LS. PLCD4 expression in the brain is elevated between 25-35 weeks post conception. Building on these observations, a loss of PLCD4 activity, for example via the stopgain mutation in PLCD4 that was identified in LSPH004 could act as an enhancer of OCRL phenotype. It is however to be noted that this variant is a heterozygous one and the biochemical impact of this variant will depend on the allelic expression pattern of PLCD4 in LSPH004 cells.

In addition to the variants in CEP290 and PLCD4, variants unique to LSPH004 were also found in ANO6, ARHGEF3 ZFP2 and POSTN. All these genes are expressed in the brain during human fetal development and may hence contribute to brain phenotypes. Likewise stop gain variants found in both LSPH002 and LSPH003 but not LSPH004 (Fig 4D) could act as suppressers of the neurodevelopment phenotype; these include the genes CAGE1, ECHCD2, MGAT4D, NCAM1, NLRX1 and SAPCD1. Some of these such as NCAM1 and NLRX1 that show unique and upregulated expression in the brain during development may be of particular importance. Lastly a set of de novo variants in genes implicated in neural function such as SCN7A, PDCD11, SORSC3, MAPK42, CDON,C5 and RGS7 may contribute to the variable phenotypes. The functional significance of these needs to be established through experimental analysis.

In summary, our work provides insights into the variable neurodevelopmental phenotype associated with LS and may provide ways to predict the evolution of the disease and hence better clinical management in individual patients.

## Acknowledgements

This work was supported by the Department of Atomic Energy, Government of India, under Project Identification No. RTI 4006. a Wellcome-DBT India Alliance Senior Fellowship (IA/S/14/2/501540) to PR, the Department of Biotechnology, Government of India and the Pratiksha Trust. PS was supported by a postdoctoral fellowship from the Science & Engineering Research Board, Government of India (DST No: PDF/2015/000310). We thank the Central Imaging Facility, Genomics Facility and High Performance Computational Facility at NCBS for support. We thank Shubra Bhattacharya for assistance.

## Materials and Methods

### DNA extraction and Sanger sequencing

Genomic DNA was isolated from the patient's peripheral blood samples following manufacturer's instruction. All 23 individual exons were amplified by PCR using Taq polymerase and oligonucleotides following the manufacturer's instruction. The PCR amplicons were cleaned up using QIAquick PCR Purification Kit and sequenced by sanger sequencing using forward or/and reverse primers. The list of primers used is presented in Supplementary Table 4

### Molecular cloning of UAS-HA::hOCRL and UAS-HA::hOCRL^688C-T^

HA::hOCRL cDNA was amplified from the pcDNA3-HA::hOCRL (Addgene-Plasmid# 22207). Not1 and Xba1 restriction sites were used to amplify the amplicon of 2730 bp and ligated into pUAST-attB vector (Drosophila Genomic Resource Center-Stock#1419). Site directed mutagenesis was used to introduce 688C-T mutation in pUAS-HA::hOCRL. Oligonucleotides used:

Not1-hOCRL-FP: GCTGCGGCCGCATGTACCCATACGACGTC
Xba1-hOCRL-RP: GCTTCTAGATTAGTCTTCTTCGCT
hOCRL688^C-T^-FP: ATCCTGGCAAAGTGAGAGAAAGAATA
hOCRL688^C-T^-RP: ATTCTTTCTCTCACTTTGCCAGGATA

### Protein expression and localization studies

The constructs were transfected in S2R^+^ cells stably expressing Actin-GAL4 using Effectene (Effectene Qiagen Kit [301425]) as per manufacturer’s protocol. Cells were processed for immunostaining using protocol as described in Panda et.al. (2018). Antibody used: mouse anti-HA(1:100, CST [2367S]). Appropriate secondary antibodies conjugated with a fluorophore were used at 1:300 dilutions [Alexa Fluor 488/568/633 IgG, (Molecular Probes)]. Cell extracts were processed for western blotting using protocol as described in (Trivedi et al., 2020). Antibody used: mouse anti-HA(1:1000, CST [2367S]) and mouse anti-Tubulin (1:4000, DHSB[E7c]. Appropriate secondary antibody conjugated to horseradish peroxidase were used at 1:10,000 dilution(Jackson Immunochemicals).

### Next generation sequencing

The Whole Genome DNA libraries were constructed using TruSeq Nano DNA LT Sample Preparation Kit Set A (24 Samples), Catalog number-FC-121-4001 and whole exome DNA libraries were prepared using Truseq Exome kit, Catalog number-FC-150-1001 according to the manufacturer’s instructions (Illumina, USA). Next generation sequencing of libraries were performed using Illumina Hiseq 2500 for 2×125 bp. The reads were trimmed off the adapter sequences using Illumina bcl2fastq2 conversion software v2.20.

### Bioinformatics Analysis

#### Sequencing reads processing & alignment

Raw sequence reads were assessed using FastQC (www.bioinformatics.babraham.ac.uk/projects/fastqc/). Paired-end raw reads with a Phred score more than Q20 were filtered using Prinseq lite v0.20.4 (Schmieder and Edwards, 2011) and were aligned to the human reference genome hg19 (GRCh37) using BWA v0.5.9 (Li and Durbin, 2009). PCR duplicates were marked using Picard (http://broadinstitute.github.io/picard/). Conversion of the sequence alignment file (SAM to BAM), indexing and sorting were done by samtools version 1.5 (Li et al., 2009). Base quality score recalibration (BQSR) and INDEL realignment was performed using Genome Analysis Tool Kit (GATK) v3.6 (Depristo et al., 2011).

#### Variant Calling

GATK HaplotypeCaller (Poplin et al., 2018) was employed for SNP and INDEL discovery. The HaplotypeCaller was performed in the GVCF mode with --emitRefConfidence parameter. Joint genotyping of each trios was done using GATK CombineGVCFs, followed by GATK Variant quality score recalibration (VQSR) (Depristo et al., 2011) and phase by transmission using GATK PhaseByTransmission (Francioli et al., 2016). Hard filtering parameters and recalibration parameters were employed according to the GATK Best Practices recommendations (Depristo et al., 2011). The filtered variants were annotated using ANNOVAR (Wang et al., 2010).

#### Identification of rare inherited variants

Annotated variants (SNP and INDEL) were subjected to a series of filtering and/or prioritizing steps to identify rare inherited variants in the affected children. Firstly, variants were tagged in accordance to their inheritance i.e. maternal, paternal or inherited from both the parents. The tagged variants were only considered for further analysis if the genotype of the variant and child were confidently ascertained (covered with at least a depth of 20 and mapping quality more than 30 in both the parents and the child) were only considered. A variant was assigned to be inherited from one parent, if the variant was identified in the child and the given parent with alternate allele depth of 8 or more and alternate allele frequency of more than 0.3, and if the alternate allele frequency was less than 0.05 in the other parent. If the variant was identified as a homozygous variant in the child and both the parents carried the variant with alternate allele depth of 8 or more and alternate allele frequency of more than 0.3, then the variant was tagged to be inherited from both the parents.

Further, the variants were filtered and categorized into three based on the type and functional relevance of the variant:

i. Likely Gene Disrupting (LGD) variants: Variants that are likely to be disrupting the translation of the gene to a functional protein – i.e., variants annotated as stopgain mutation, frameshift insertion, or frameshift deletion.
ii. Missense Damaging (MisDm) variants: Non-synonymous variants that were predicted by both SIFT (Kumar et al., 2009) and PolyPhen2 (Adzhubei et al., 2013) to have a deleterious effect.
iii. Missense variants: All non-synonymous variants identified.

The filtered variants were also tagged if their clinical significance was known. This was done by filtering the variants with the following annotation terms from ClinVar (Landrum et al., 2018) - Pathogenic, Likely_pathogenic, risk_factor, association, or protective. To identify rare variants, a population frequency filter of 0.01 from ExAC-SAS (South Asian samples) (Karczewski et al., 2020; Lek et al., 2016) was used and variants from the above three categories were filtered using this. Finally, only the filtered variants identified by both whole genome and exome sequencing were taken for further analysis to ensure a high-confidence variant call set.

#### Identification of de novo variants

To identify *de novo* variant candidates and reduce errors, we derived empirically validated filters from 5 studies involving large-scale analysis of *de novo* mutations in trios (Fromer et al., 2014; Jiang et al., 2013; Willsey et al., 2017; Wright et al., 2015)(Sanders et al., 2012). The annotated set of variants obtained from the GATK pipeline detailed in the Variant Calling section was further employed to identify *de novo* variants in the three OCRL probands. The following are the six criteria that were used to filter *de novo* variants:

i. The variant is identified as heterozygous in the child with alternate frequencies between 0.3 and 0.7
ii. Not identified in both the parents (alternate frequency less than 0.05)
iii. Sequencing depth of the position of the variant is at least 20 in child and the parents
iv. The alternate allele is supported by at least 8 reads in the child
v. Minimum mapping quality of 30 at that genomic position
vi. Minimum Phred-scaled genotype likelihood of at least 20

For X and Y chromosomes, the criteria were modified as:

i. The variant is identified in the child with alternate frequency above 0.3
ii. Not identified in the parent (alternate frequency less than 0.05)
iii. Sequencing depth at the position of the variant is at least 10
iv. The alternate allele is supported by at least 4 reads in the child
v. Minimum mapping quality of 30 at that position
vi. Minimum Phred-scaled genotype likelihood of at least 20

Further, the candidate *de novo* variants were filtered to remove common variants in the population (frequency filter of 0.01 in ExAC-SAS) (Karczewski et al., 2020; Lek et al., 2016). Finally, only the filtered *de novo* candidate variants identified by both whole genome and exome sequencing were taken for further analysis.

#### Curation and generation of 3-Tiered System of categorization of candidate genes

To understand the mutational burden and its relevance to the LS phenotype, a 3-tiered system of categorization of candidate genes was developed. Tier 1 genes were defined as those genes that are known for causing abnormalities, disorders or phenotypes that are diagnosed in LS or are very similar to the phenotypes of the syndrome. To curate this, phenotype terms that are diagnosed in LS were extracted from literature and categorized into three - ocular, cerebral and renal, based on the tissue that’s affected by the given phenotype. This list of phenotype terms was manually curated by an expert clinician to make sure only phenotypes relevant to LS were included. For each phenotype term, a search query was made in the OMIM dataset (Online Mendelian Inheritance in Man; https://omim.org/). If there was a match between the name or the alternative name of the OMIM phenotype and if the molecular basis of the same was known, then these OMIM entries with the causative information were extracted. The same process was repeated with the database DECIPHER (Firth et al., 2009) and repetitive entries were removed. Also, direct interactors of the gene OCRL were extracted from literature and added to this list. Hence, tier 1 essentially contained four sub-categories of set of genes – Ocular, Cerebral, Renal and OCRL Interactome. Tier 2 set of genes were defined as those genes in biological pathways where OCRL gene is critically involved. In other words, Tier 2 is a list of genes from all biological pathways regulated by OCRL. Every biological process in which OCRL gene is involved was extracted from literature and GeneCards annotation (Stelzer et al., 2016). GO terms for these biological processes were extracted and they were queried in the Gene Ontology database (Ashburner et al., 2000; Carbon et al., 2021). All the genes that are annotated with the GO term of these biological processes were extracted. These extracted gene entries were treated as Tier 2 set of genes. Tier 3 list of genes were defined as those that interact directly with tier 1 genes. For every gene in the tier 1 set of genes, a query was made in the BioGRID database (Oughtred et al., 2021). These genes were annotated with information in the BioGRID dataset to obtain the tier 1 interactor, type of experiment and literature evidence showing the interaction. The curated set of 3-tiered candidate genes are provided in the Supplementary Table 2.

#### Software and custom scripts

Variants filtering, prioritization and 3-tiered candidate genes curation were performed using in-house custom python and bash scripts. Python v2.7 or higher was used. Data visualizations were done with R v3.3.2 or higher. Variant summary plots were created using ComplexHeatmap package (Gu et al., 2016). Data representation was also performed using GraphPad Prism v8.0.0 (GraphPad Software, San Diego, California USA, www.graphpad.com).

### Ethics Approval

This work was carried out under the ethics approval provided by the Institutional Ethics Committee, St. John’s Medical College & Hospital, Bangalore (IEC Study Ref. No. 28 / 2017) and the Institutional Ethics Committee, National Centre for Biological Sciences, Bangalore (NCBS/IEC-8/002).

## Supplementary Data

**Supplementary Figure S1:**
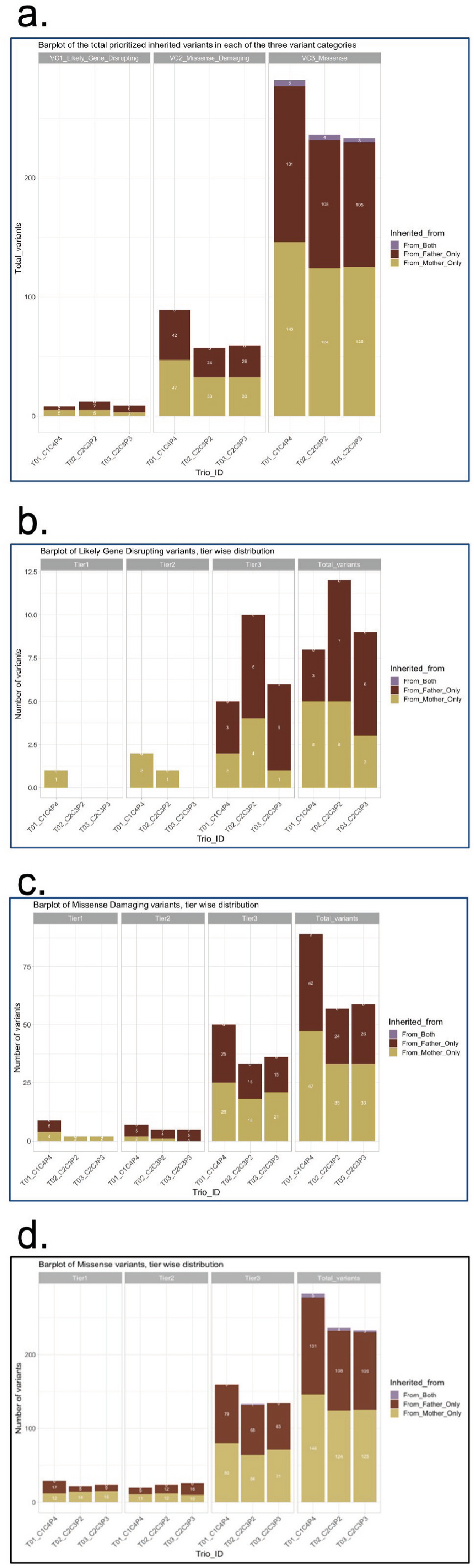
Distribution of prioritized rare inherited variants: (a) The number of rare inherited variants that were prioritized in each patient. The prioritized rare inherited variants are categorized into three (Likely Gene Disrupting, Missense Damaging and Missense) and the distribution for each category is shown. The bar plots are stacked to show the distribution of variants that were inherited from father, mother or both. (b) The prioritized variants were annotated with the 3-tiered system of categorization of candidate genes. The tier-wise distribution for the prioritized rare inherited variants is shown for Likely Gene Disrupting (LGD) variants, (c) Missense Damaging (MisDm) variants and (d) Missense (Mis) variants are shown in stacked bar plots. The IDs T01_C1C4P4, T02_C2C3P2, and T03_C2C3P3 correspond to the patients LSPH004, LSPH002 and LSPH003 respectively.

**Supplementary Table 1:** List of prioritized rare inherited and de novo variant, along with relevant annotations. Sheet one of the excel file in a n information sheet describing the contents of all other sheets.

**Supplementary Table 2:** Analysis of individual clinical features in LS patients genotyped as carrying a stop codon prior to the start of the 5’ phosphatase domain. Individual phenotypes where noted are marked with (y) and not noted (n). Note determined is n.d. Individual studies from which this data are collated are listed. Graph shows the distribution of known LS phenotypes in a cohort of patients all of whom carry a stop codon mutation prior to the start of the 5’phosphatase domain. Y-axis shows the individual clinical features. X-axis in the % of patients in this cohort (n=12) who display a particular phenotype.

**Supplementary Table 3:** Gene lists from the three-tiered categorization (3TS) of candidate genes for OCRL. Sheet 1 in an information sheet. Additional annotation in each list is provided, as relevant.

**Supplementary Table 4:** List of primers used for the Sanger sequencing of the OCRL gene in patient samples.

