## supplemental table 4 for "Genomic sequencing of Lowe syndrome trios reveal a mechanism for the heterogeneity of neurodevelopmental phenotypes"

**Supplementary Table 4: Primers used for sanger sequencing:**

| Exon | Oligonucleotide | Sequence (5'-3') |
|---|---|---|
| 1 | F1 | GCTGGACCTGGTAGAGACGCCCCC |
| | R1 | CCACCCGGTTGCCCCGCCCC |
| 2 | F2 | CTTCGCCCCGCAGTGCACCACT |
| | R2 | CGTGGCTTCAGTAGGAGGAC |
| 3 | F3 | CCTAGAGCAATATCTAGCTGTCA |
| | R3 | TACATAATGCAGTAATGACCGAG |
| 4 | F4 | TGAGGAGTTCCATTTGGTTAC |
| | R4 | TTCTTAGGCTTAGCCTACATG |
| 5 | F5 | GTGTGCTTCCTATATTAGAGAT |
| | R5 | ATCAGCTATCAGGGCTTAAAG |
| 6 | F6 | CTGTATCCAAAAATGGTGCTGG |
| | R6 | AGGAGAATGTGTCTGACATCAG |
| 7 | F7 | ACCACTGATCAAATTGTGATC |
| | R7 | ACTATAATTCTCCTACAAGGC |
| 8 | F8 | GCCTTGTAGGAGAATTATAGT |
| | R8 | CTTAAATATCAACAGGCCACT |
| 9 | F9 | CATACCTTTGTATGGAAGCG |
| | R9 | TACAGTAGGTTTTACCAACAGT |
| 10 | F10 | GTGAACAGAGCAGTTCTATAA |
| | R10 | GATAATGGAATAACTCCCGGT |
| 11 | F11 | ACTAGTATATCATCTTGATGGA |
| | R11 | GCAACTGAACTTTACATGAAC |
| 12 | F12 | GTTCATGTAAAGTTCAGTTGC |
| | R12 | ATTTAATCTCTACACTATCCA |
| 13 | F13 | AGTGGTGAGTGAGCCCTTAT |
| | R13 | CAGTAAGACGTTTCCATCACTCC |
| 14 | F14 | ATAGGAACAGTGGCTTATCAAC |
| | R14 | CAATATGGAGGGTCAGAAAT |
| 15 | F15 | TAACAAGAGAGCCTAACCCT |
| | R15 | CCTCTAGTAATTGATACTTAACAC |
| 16 | F16 | GTTAGATAATCTCCAAGGGAG |
| | R16 | TTCTGTGCTAACCACAGTGAG |
| 17 | F17 | ATCCTCTATGGAATAATCCAAC |
| | R17 | TCATGACATCACCAGCAG |
| 18 | F18 | GCTTTCCCACTGGAGGTTTTCC |
| | R18 | AGGACGTCACTTAAGTATTGAG |
| 19 | F19 | GCATGACCAGAATTTGAAGGA |
| | R19 | GAGGTGTTGTGATTTCCTATAG |
| 20 | F20 | GGTAATCATCATAACCTCAG |

| | | |
|---|---|---|
| | R20 | TATAGGTGCGAAGAAGAGT |
| 21 | F21 | AGCTTGAGGAAAGGAGCTC |
| | R21 | CCTATCTGGCTCTGTAAAACT |
| 22 | F22 | GGGTCCTGCAAGGGTTGG |
| | R22 | GGGGACAAGGACTATTGACTGC |
| 23 | F23 | GGCAGTCAATAGTCCTTGTC |
| | R23 | GACCCTTCCTGTGGCATGA |
